# Deletion of the *MALAT1* RNA 3’ end Promotes Transcript Decay and Inhibits Proliferation in Gastric and Breast Cancer Cells

**DOI:** 10.64898/2026.05.20.726613

**Authors:** Aramis Cortes-Arias, Valeria Valdes, Marcelo Muñoz-González, Diego Leiva, Andrew Acevedo, Michelle Tobar-Lara, Nicole Farfán, Lilly Oni, Eduardo Quiñones-Medina, Veronica A. Burzio, Adriana Rojas, Roberto Munita, Srinivas Somarowthu, Fernando J. Bustos, Rodrigo Aguilar

## Abstract

The long non-coding RNA *MALAT1* is a conserved oncogenic driver whose function relies on a 3’ triple-helix motif. While its biochemistry is well-characterized *in vitro*, the endogenous requirement for this motif in regulating the stability of the transcript and other genes residing in its locus remains unclear. In this study, we employed a dual-sgRNA CRISPR-Cas9 approach to systematically excise triple-helix-forming sequences from the native *MALAT1* locus in gastric (AGS) and breast (MCF7) cancer cells. Our findings demonstrate that the 3’ end strongly contributes to *MALAT1* stability. Perturbations ranging from genomic deletions to a single-base changes trigger transcript collapse and rapid exonucleolytic decay, while the biogenesis of the small RNA mascRNA (a byproduct of *MALAT1*, also involved in cancer) remains decoupled and unaffected. *In cellulo,* DMS probing reveals that edited transcripts retain structural complexity in the 3’ region. Phenotypically, structural disruption of the 3’ end significantly impairs proliferation of both cancer cellular models. These results identify the 3’ triple-helix as a determinant of *MALAT1* stability and provide endogenous validation for its role in the analyzed AGS and MCF7 cells.

## Introduction

The long non-coding RNA (lncRNA) Metastasis-Associated Lung Adenocarcinoma Transcript 1 (*MALAT1*) has emerged as a central regulator in oncology [1]. Since its initial discovery in patients with non-small cell lung cancer [2], its over-expression has been documented across a wide spectrum of malignancies, including breast [3–5] and gastric cancer [6], where it acts as a driver of cell proliferation, migration, and invasion (a comprehensive list of *MALAT1*-related finding in different cancer types and their associated cell lines can be found at [7]). Due to its high conservation and consistent upregulation in solid tumors, *MALAT1* is frequently cited as a primary candidate for RNA-targeted therapeutic intervention [1].

At the molecular level, the 7,472-nt *MALAT1* transcript is primarily accumulated within nuclear speckles [8], serving as a structural scaffold for pre-mRNA splicing factors and modulating the distribution of serine/arginine (SR) proteins [8]. Unlike most Polymerase II-transcribed RNAs, the stability and nuclear accumulation of *MALAT1* are not maintained by a canonical poly(A) tail. Instead, its longevity is sustained by an Expression and Nuclear Retention Element (ENE) located at its 3’ terminus [9]. During *MALAT1* biogenesis, this ENE facilitates the formation of a highly conserved triple-helix structure, characterized by the interaction between two U-rich internal tracts and a genome-encoded, short A-rich sequence [9]. This unique structural motif protects the transcript from 3’-5’ exonucleolytic decay, bypassing the need for a standard polyadenylation signal, making it a valuable module for engineering chimeric transcripts and aptamers with increased half-lives and stability [10, 11].

The maturation of the *MALAT1* 3’ end is a complex process involving the cleavage of the primary transcript by RNase P and RNAse Z, which releases a short tRNA-like byproduct named mascRNA (*MALAT1*-associated small cytoplasmic RNA)[12]. While *MALAT1* remains in the nucleus, mascRNA is exported to the cytoplasm, where it has been implicated in cancer progression by modulating cellular metabolism [13] and serving as a molecular scaffold for protein stabilization [12]. Furthermore, report indicate that *MALAT1* locus harbors a transcriptionally active antisense partner, *TALAM1*, which spans approximately 8,100 bases and that is expressed at lower levels than *MALAT1* in breast cancer cells [14, 15].

While the correlation between the ENE and *MALAT1* stability is well-supported by biophysical assays [9, 16] and the use of small molecules [17], these models often fail to account for the complex interplay between transcription and processing within the endogenous genomic environment. Recently, using CRISPR-Cas9 technology, it was demonstrated that a single-guide RNA targeting the vicinity of the ENE to introduce indels could reduce *MALAT1* expression by approximately 50% in lung cancer cell lines [18]. However, a precise dissection of how the specific triple-helix-forming sequences regulate the *MALAT1* locus, including its associated transcripts mascRNA and *TALAM1*, remains to be fully characterized in their native chromosomal context.

In this study, we employed a dual-sgRNA CRISPR-Cas9 approach to systematically excise triple-helix-forming sequences from the endogenous *MALAT1* locus in both gastric (AGS) and breast (MCF7) cancer cell lines, models where the *MALAT1* contribution to proliferation has been documented [7, 19, 20]. We characterized the downstream effects of this structural deletion on transcript stability, the expression of mascRNA and *TALAM1*, and the resulting impact on cancer cell proliferation. Additionally, using *in cellulo* DMS probing, we evaluated whether these modifications produced a structured RNA towards the 3’ end. Our findings demonstrate that the 3’ triple-helix is a critical determinant of *MALAT1* stability, impacting cell proliferation in MCF7 and AGS cancer models.

## Materials and Methods

### Cell culture

All cell lines were maintained in a humidified atmosphere at 37°C with 5% CO_2_. The gastric cancer cell line AGS (ATCC #CRL-1739) [21] and its CRISPR-derived variants were cultured in F-12K medium. The breast cancer cell line MCF7 (ATCC #HTB-22) [22] and its derivatives were maintained in DMEM medium. All culture media were supplemented with 10% (v/v) Fetal Bovine Serum (FBS) and 100 U/mL Penicillin/Streptomycin. All cell culture reagents were purchased from Thermo Fisher Scientific (USA).

### CRISPR-Cas9-Mediated Genomic Deletions

Single-guide RNAs (sgRNAs) targeting the *MALAT1* 3’ end were designed using the Benchling platform [23]. Synthetic oligos (**Table 1**) were cloned into the pL-CRISPR.EFS.tRFP plasmid (plasmid was a gift from Benjamin Ebert, Addgene plasmid #57819; http://n2t.net/addgene:57819; RRID:Addgene_57819 [24]). To generate genomic deletions, cells were co-transfected with plasmid combinations (p6042+p6048, p6042+p6087, or p6048+p6087) using Lipofectamine 3000 (Thermo Fisher Scientific) according to the manufacturer’s instructions. A pL-CRISPR.EFS.tRFP vector lacking sgRNA inserts (“RFP empty”) served as a negative control. 72 h post-transfection, RFP-positive cells were isolated via fluorescence-activated cell sorting (FACS) using a FACSAria™ III Cell Sorter (BD Biosciences, USA) located at Universidad Andres Bello. Single-cell clones were subsequently established through limiting dilution to ensure genomic homogeneity. For each targeted deletion, three independent clones per cell line were selected for expanded downstream analysis.

**Table 1.**
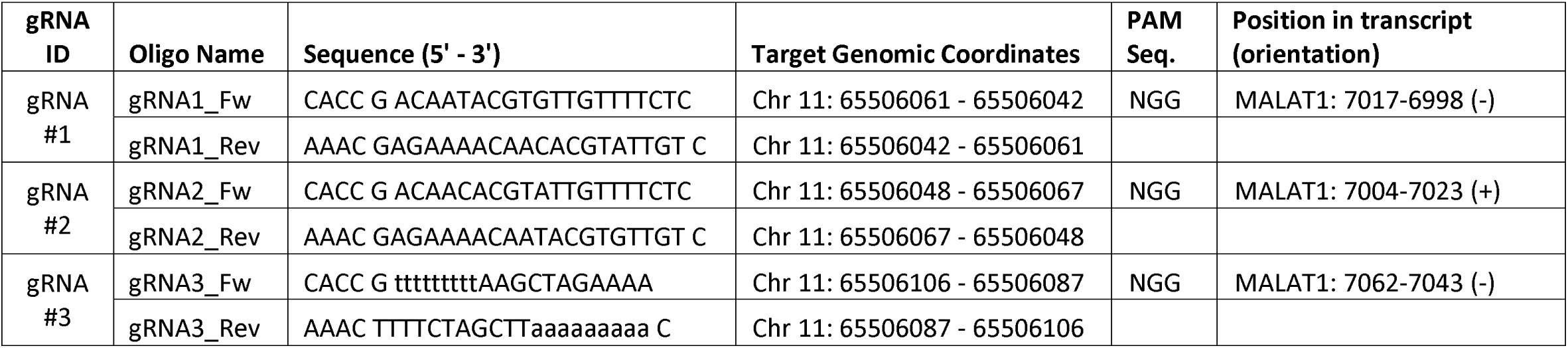
Synthetic oligonucleotides used for gRNA-encoding plasmid construction.

### In Silico Off-Target Prediction

Single-guide RNAs (sgRNAs) were designed to target the conserved 3⍰ triple-helix domain of the *MALAT1* locus using Benchling. Benchling, CCTop, and CRISPR-off were used for off-target predictions. Because targeting was strictly constrained to this functional structural motif, the available sequence window and SpCas9 NGG PAM sites were severely limited, precluding the selection of alternative sgRNAs with lower off-target risk. *In silico* off-target analysis identified the following potential off-target profiles: sgRNA 1 and 2 each displayed 2 potential off-target sites with 2 mismatches, and 13 with 3 mismatches; sgRNA 3 displayed 33 potential off-target sites with 2 mismatches, and 311 with 3 mismatches. Targeted sequencing of predicted off-target loci was not performed.

### Molecular Characterization of CRISPR Clones

To confirm the targeted genomic deletions, total DNA was extracted and analyzed by conventional PCR using the KAPA2G Robust HotStart PCR Kit (Merck, Germany). Amplicons were generated using primers flanking the deletion sites (Forward: 5’-GGTGGGAATGCAAAAATTCTCTG-3’; Reverse: 5’-ACAGAAAGAGTCCTGAAGACAGAT-3’; primers from IDT, USA) and resolved on 1% (w/v) agarose gels stained with GelRed (Biotium, USA). Targeted bands were excised and purified using the Zymoclean Gel DNA Recovery Kit (Zymo Research, USA). To characterize the specific repair junctions, purified PCR products were subcloned into the TOPO® TA Cloning® Kit (Thermo Fisher Scientific) and subjected to Sanger sequencing (Macrogen, Chile) using primers targeting SP6 and T7 universal promoters. At least 2 colonies per clone were sequenced to verify allelic composition

### RNA extraction

Total RNA was isolated from cells preserved in TRIzol reagent (Thermo Fisher Scientific) using the Direct-zol RNA Kit (Zymo Research), and incorporating the on-column DNase I treatment to eliminate genomic DNA contamination. RNA concentration and purity were assessed using a NanoDrop spectrophotometer (Thermo Fisher Scientific).

### Reverse Transcription (RT), Strand-Specific RT, and Quantitative PCR

For gene expression analysis, 1 μg of total RNA was reverse-transcribed using SuperScript IV RT (Thermo Fisher Scientific). General quantification was performed using random hexamers, while strand-specific detection of *TALAM1* was achieved using 1 pmol of a tagged antisense-specific primers per 20 μL reaction. The resulting cDNA was diluted to a final volume of 100 μL. Quantitative real-time PCR (qPCR) was performed using Brilliant II SYBR Green QPCR Master Mix (Agilent Technologies) on a Quantstudio I Thermal cycler (Thermo Fisher Scientific). Relative expression levels were calculated using the 2^-ΔΔCt^ method. Primers are listed in **Table 2**

**Table 2.**
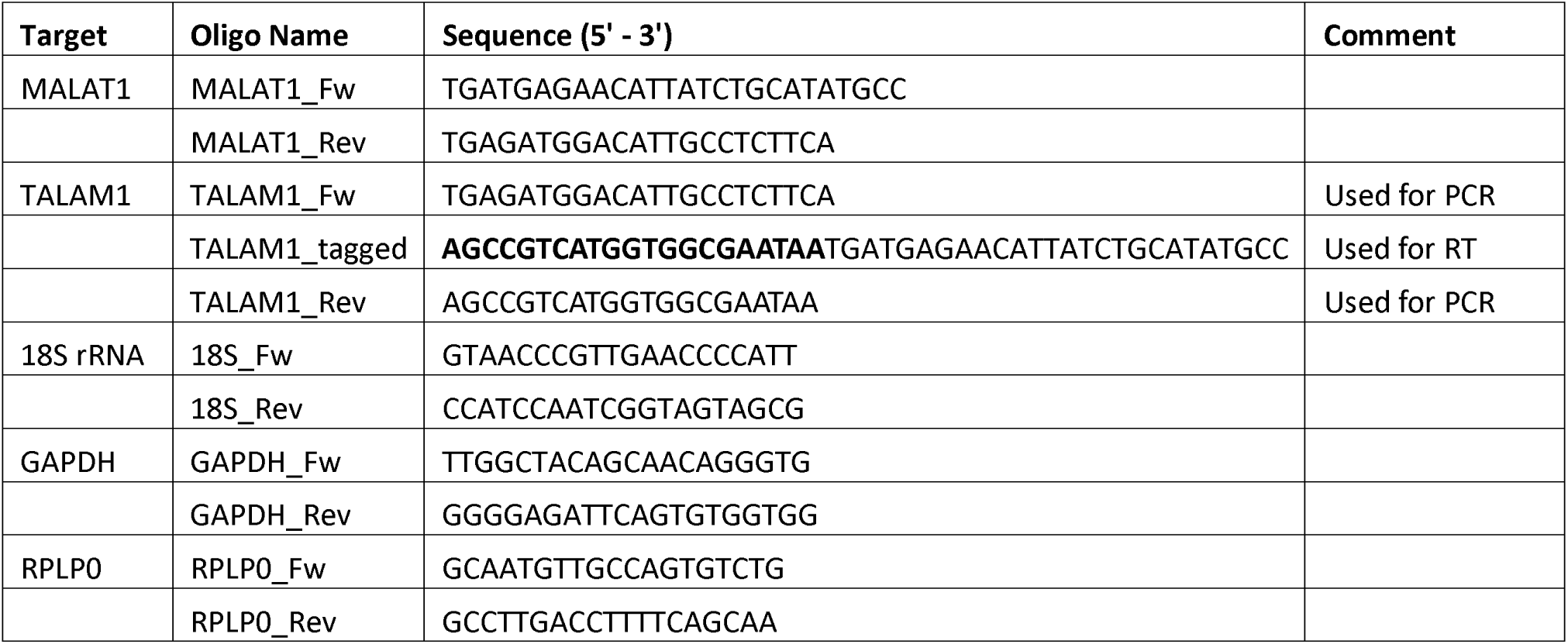
Synthetic oligonucleotides used for quantitative and digital PCR.

### Digital PCR (dPCR)

dPCR was performed using the QIAcuity EvaGreen PCR Kit (QIAGEN) on a QIAcuity Digital PCR System (QIAGEN). Amplification was carried out using 0.4 μM of specific primers and 3 μL of cDNA template in a final reaction volume of 12 μL, which was partitioned into a Nanoplate 8.5K 24-well (QIAGEN). Absolute quantification was determined using the QIAcuity Software Suite version 3.1.0.0. Primers are listed in **Table 2**.

### RNA Fluorescence *In Situ* Hybridization (RNA-FISH)

To visualize the subcellular localization of *MALAT1*, MCF7 and AGS cells were grown on glass coverslips, fixed with 3.7% (v/v) formaldehyde (Sigma-Aldrich, USA) for 10 minutes at room temperature, and permeabilized in 70% (v/v) ethanol (Merck) at 4°C for at least 1 hour. *MALAT1* transcripts were detected using Stellaris® FISH Probes (Biosearch Technologies, USA) labeled with Quasar® 570 Dye (Cat. #SMF-2035-1), consisting of a pooled set of oligonucleotides tiling the full-length transcript. Total mRNAs were detected using Stellaris® Positive Control T30-FAM (Cat. #T30-FAM-1). Hybridization was performed at 37°C for 16 hours in a humidified chamber, following manufacturer’s instructions. Post-hybridization, cells were washed with Stellaris Wash Buffers A and B to remove non-specific binding. Nuclei were counterstained with NucBlue™ Fixed Cell ReadyProbes™ Reagent (DAPI) (Thermo Fisher Scientific). Coverslips were mounted using Fluoromount-G (EMS) and imaged using a Nikon epifluoresence microscope (Japan) controlled by NIS-Elements software. Images were processed and analyzed using FIJI software [25], maintaining consistent acquisition settings across all experimental groups.

### RNA Stability Assays (Actinomycin D Treatment)

To determine *MALAT1* and *TALAM1* transcript half-life, MCF7 and AGS cells were treated with 1 μg/mL Actinomycin D (Cayman Chemical, USA) to inhibit *de novo* transcription. Total RNA was harvested at t = 0, 3, 6, and 10 h post-treatment. RNA decay kinetics were determined by plotting the remaining RNA fraction over time. The transcript half-life (t_1/2_) was determined by non-linear regression using a one-phase exponential decay model, with the initial plateau (Y_0_) constrained to 1.0

### *In Cellulo* DMS Probing

#### Chemical probing of cellular MALAT1 RNA

AGS cells were seeded in 10 cm plates 72 h before chemical probing or until they reached 70– 80% confluency. After one wash with DPBS, cells were resuspended in DPBS supplemented with 80 U RNase inhibitor. RNA structure probing was performed using Dimethyl sulfate (DMS)[26]. A final concentration of 0.6% DMS was added to treated cells, while an equal volume of pure ethanol was added to control cells. Cells were incubated for 6 min at 37 °C, then rapidly transferred to 4 °C, and the reaction was quenched by adding a volume of quenching solution (1.43 M β-mercaptoethanol) equal to the volume of DMS added. Cells were washed once with quenching solution at 4 °C and incubated in TRIzol Reagent (Thermo Fisher Scientific) for 5 min at room temperature. Total RNA was then extracted using the Direct-zol RNA Miniprep Kit.

#### Reverse transcription of modified RNA

For *in vivo* structural probing, reverse transcription was performed using MarathonRT [27]. A total of 1 μg RNA was mixed with 1 μL of 2 μM gene-specific primers. The mixture was incubated at 75 °C for 2 min and immediately placed on ice. Next, 8 μL of 2.5X RT buffer (125 mM Tris-HCl, pH 7.5, 500 mM KCl, 12.5 mM DTT, 1.25 mM dNTP, 6.25 mM MnCl2), 4 μL of 100% glycerol, and 10 U MarathonRT were added to a final reaction volume of 20 μL. Samples were incubated at 42 °C for 3 h, and the reverse transcriptase was inactivated by incubation at 75 °C for 10 min. Following reverse transcription, cDNAs were purified using G-25 spin columns according to the manufacturer’s instructions.

#### Amplicon Sequencing

cDNA was PCR-amplified using specific primers (Set 1: Malat1_6335_Fwd: 5’ CCCCTGGGCTTCTCTTAACAT 3’, Malat1_6819_Rev: 5’ ACATGTTCCCACCCAGCATTA 3’; Set 2: Malat1_6626_Fwd: 5’ TGGTGTCGAGGTCTTTGGTG 3’, Malat1_7232_Rev: 5’ GTGCTGTTACCTCCCACCTC 3’) using Q5 Hot Start DNA Polymerase (NEB). The number of PCR cycles was limited to 30 or fewer. Samples were then purified using the DNA Clean & Concentrator and quantified using the Qubit 4 Fluorometer. A total of 320 ng DNA from each sample was submitted for long-read dsDNA amplicon sequencing (Plasmidsaurus, KY, USA).

#### In-cell DMS data analysis

DMS mutational profiling analysis of the MALAT1 amplicons were performed using RNAFramework (v2.9.6) [28]. The workflow included the following tools: rf-map, rf-count, and rf-index for read processing and mutation counting, followed by rf-norm for normalization using the following parameters: -sm 3, -nm 2, -rb AC, and -norm-independent. Structure prediction was performed using rf-fold with in-cell DMS reactivities as constraints. In-cell DMS reactivities from two replicates were used. Mutation rates were calculated by comparing A and C mutation frequencies between control and DMS-treated samples within RNA count (.rc) files generated after rf-count processing, and the resulting mutation counts were displayed in mutation graphs.

### Northern Blot Analysis of Small RNAs

To detect the expression of small RNAs (mascRNA and *U6*), total RNA was extracted from AGS and MCF-7 cells using TRIzol reagent. For Northern blot analysis, 3.5-7 μg of total RNA were denatured in loading buffer (95% formamide, 18 mM EDTA, 0.025% SDS, and 0.1% bromophenol blue) at 85°C for 10 minutes and immediately chilled on ice. Samples were resolved on a 15% (w/v) polyacrylamide gel containing 7 M urea in 1X TBE buffer. Electrophoresis was performed at 200V in 0.5X TBE buffer until the bromophenol blue front reached 1 cm from the bottom of the gel. RNA was transferred onto a Hybond N+ nylon membrane (GE Healthcare) via electroblotting at 250 mA for 1 hour at 4°C in 0.5X TBE. Following transfer, the RNA was immobilized by UV crosslinking at 1500 mJ/cm². Membranes were pre-hybridized for 30 minutes at 40°C in a solution containing 5X SSC, 50% formamide, 0.1% SDS, 3X Denhardt’s solution, and 200 μg/mL sheared salmon sperm DNA. Hybridization was performed overnight at 45°C with specific biotinylated oligonucleotide probes synthesized by IDT for mascRNA (5’-AGACGCCGCAGGGATTTGAACCCCGTCCTGGAAACCAGGAGTGCCAACCACCAGCATC/3Bio/-3’; 150-300 pmol/mL) and U6 snRNA (5’-GAACGCTTCACGAATTTGCGTGTC/3Bio/-3’; 50 pmol/mL). Sequential stringency washes were performed at 40°C: 15 minutes in 2X SSC/0.1% SDS, 15 minutes in 1X SSC/0.1% SDS, and 15 minutes in 0.5X SSC. Chemiluminescent signals were detected using the Chemiluminescent Nucleic Acid Detection Module (Thermo Fisher Scientific) and visualized with by 12-minute exposure on a C-DiGit Blot Scanner (LI-COR Biotech, Model 3600). Densitometric analyses were performed using Li-cor image studio v3.1.4.

### Cell proliferation study

For growth curve analysis, cells were seeded into 12-well plates at an initial density of 1.0×10^5^ cells per well. Cell proliferation was monitored every 24 hours for a total of 72 hours. At each time point, cells were harvested by trypsinization and total cell counts were determined using a Luna-III™ Automated Cell Counter (Logos Biosystems, South Korea). To distinguish live from dead cells, samples were diluted 1:1 with 0.4% (w/v) Trypan Blue solution (Invitrogen). All experiments were performed in three independent biological replicates

### Flow cytometry

Cell apoptosis and necrosis were evaluated using Annexin V and Propidium Iodide (PI) staining, respectively. Briefly, cells were harvested, washed in cold phosphate-buffered saline (PBS), and centrifuged. The supernatants were discarded, and the cell pellets were resuspended in annexin-binding buffer (10 mM HEPES, 140 mM NaCl, and 2.5 mM CaCl_2_, pH 7.4). For each experimental condition, 300,000 cells were resuspended in a final volume of 300 µL of Annexin-binding buffer. Cells were then stained by adding 12 µL of Annexin V conjugate (Thermo Fisher Scientific; equivalent to 4 µL per 100,000 cells) and Propidium Iodide at a final concentration of 40 µg/mL (Sigma-Aldrich). Samples were mixed gently and incubated for 15 minutes at room temperature in the dark.

Immediately following incubation, the stained cells were analyzed by flow cytometry using a FACSAria III cytometer (BD Biosciences, USA). Fluorescence excitation was performed using a 488 nm laser. Annexin V emission was detected at 530 nm, and PI fluorescence was detected at 610 nm. The cell populations were distinguished as viable cells (low fluorescence), early apoptotic cells (high Annexin V fluorescence, low PI), and necrotic or late apoptotic cells (double positive for Annexin V and PI). Data analysis was performed from a minimum of 30,000 acquired events per sample.

### Statistical analysis

All experiments were performed in at least three independent biological replicates, and data are presented as the mean ± standard deviation (SD) unless otherwise specified. Statistical significance between two groups (e.g., control vs. single CRISPR clone) was determined using an unpaired Student’s *t*-test. For comparisons involving multiple groups (e.g., multiple CRISPR clones vs. RFP empty), a one-way analysis of variance (ANOVA) followed by Tukey’s post-hoc test was applied. Proportional data from cell proliferation growth curves and RNA decay assays were analyzed using two-way ANOVA to evaluate the interaction between time and genotype. RNA half-lives (t_1/2_) were calculated by fitting a non-linear one-phase exponential decay model (Y=(Y_0_−Plateau)⋅e^−K⋅X^ + Plateau). All statistical analyses and curve fittings were performed using GraphPad Prism 9 (GraphPad Software, USA). P-values less than 0.05 were considered statistically significant.

## Results

### Genomic Deletion of the *MALAT1* 3’ Triple-Helix Motifs Reduces Transcript Levels in Gastric and Breast Cancer Cell Lines

To investigate the endogenous role of the 3’ triple-helix in *MALAT1*, we employed a dual-sgRNA CRISPR-Cas9 approach to excise specific triple-helix-forming sequences within the ENE motif. To ensure that our findings were not dependent on a specific cellular lineage, we used two distinct human cancer models: the gastric cancer line AGS and the breast cancer line MCF7, both known to overexpress *MALAT1* [7, 20, 29].

Our CRISPR strategy utilized three combinations of sgRNAs designed to truncate triple-helix-forming elements of the ENE (Figure 1A). Following transfections, cell pooling, and FACS-based enrichment of RFP^+^ populations (Figure 1B, Supplementary Figure S1), single-cell clones were established via limiting dilution. In AGS cells, PCR screening of 13 monoclones revealed that while the majority retained the wild-type (WT) 300-bp amplicon, clones #1, #3, and #5 exhibited a single truncated band of lower size, indicating biallelic deletions (Figure 1C, Supplementary Figure S2). Subsequent TOPO-TA cloning and Sanger sequencing characterized the precise nature of these genomic excisions: clone #1 harbored a 42-bp deletion (removing the first U-rich motif and the conserved loop); clone #3 exhibited a 52-bp deletion (excising the distal nucleotide of the first U-rich motif, the conserved loop, and the second U-rich motif); and clone #5 featured a 82-bp deletion (extending from 43 bp upstream of U-rich motif 1 through part of the conserved loop) (Figure 1D, Supplementary Figure S3). In sum, we successfully deleted essential triple-helix-forming sequences in the AGS genomic context.

**Figure 1.**
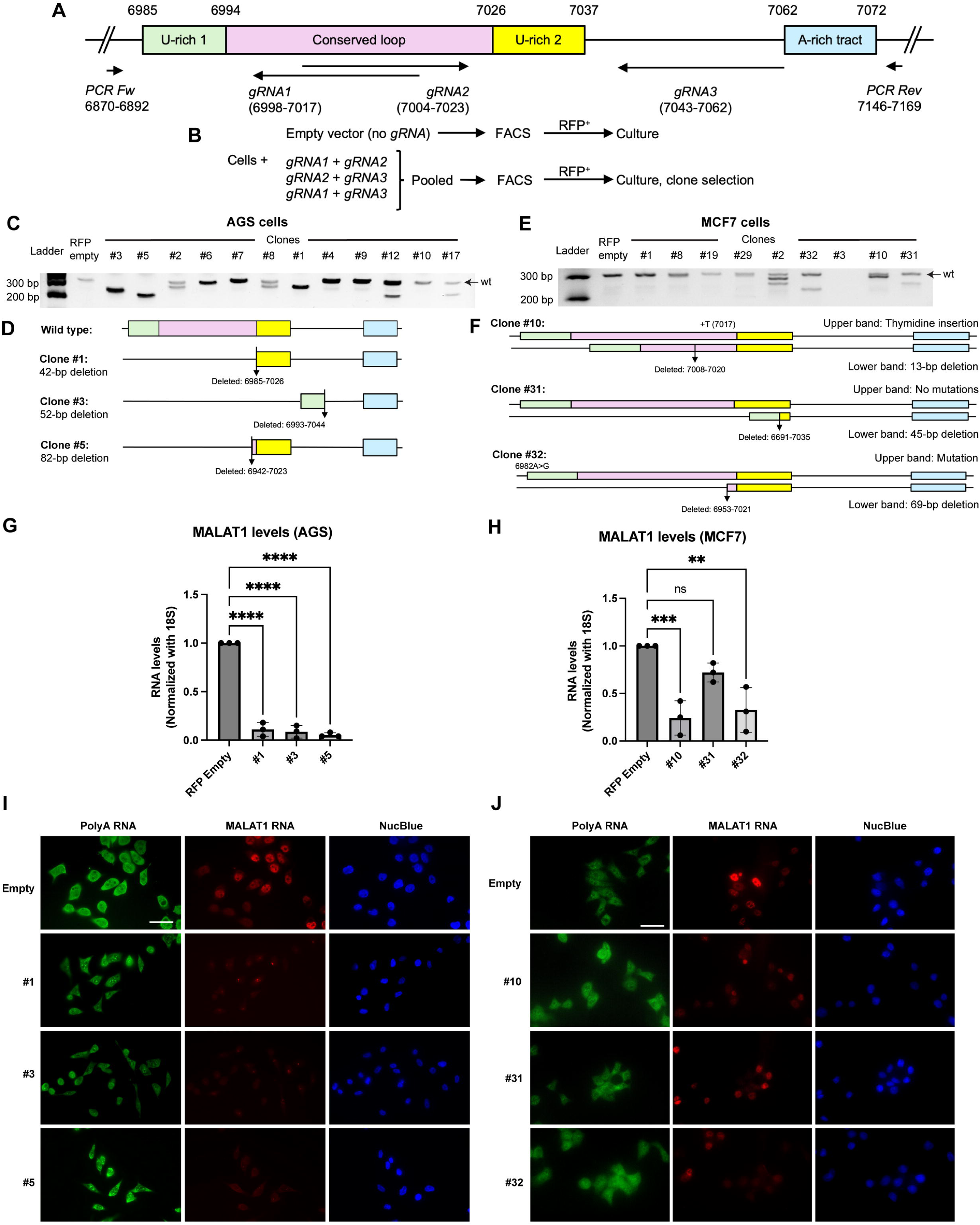
CRISPR-Cas9-mediated deletion of the MALAT1 3’ triple-helix motifs leads to reduced MALAT1 transcript levels. (A) Schematic representation of the *MALAT1* 3’ end locus and the dual-sgRNA strategy targeting the ENE motifs. Numbers correspond to the position of guide RNAs (gRNAs) and PCR primers considering nucleotide positions within the transcript. (B) Summary of the experimental approach, including FACS enrichment of RFP-positive cells. (C) PCR screening of AGS single-cell clones; truncated amplicons in clones #1, #3, and #5 indicate biallelic deletions. (D) Sequence alignment of the repair junctions for AGS clones #1, #3, and #5 compared to the wild-type (WT) sequence, highlighting the excised triple-helix motifs. (E) PCR screening of MCF7 clones showing monoallelic deletions in clones #10, #31 and #32. (F) Schematic of the heterozygous modifications in MCF7 clones. (G–H) RT-qPCR quantification of steady-state *MALAT1* levels in AGS (G) and MCF7 (H) edited clones relative to controls. Data represent mean ± SD of three biological replicates; ∗∗∗∗: *Padj*<0.0001, ∗∗∗: *Padj*=0.0009, ∗∗: *Padj*=0.0020 by one-way ANOVA. (I–J) Representative RNA-FISH images of three biological replicates using tiling probes against full-length *MALAT1* in AGS (I) and MCF7 (J) cells. Nuclei were counterstained with DAPI and total mRNA was detected with a poly-T probe. Scale bars = 20 μm

In contrast to the AGS model, screening of 35 MCF7 clones yielded exclusively monoallelic deletions, with clones #10, #31 and #32 exhibiting both wild-type-sized (WT) and truncated amplicons (Figure 1E, Supplementary Figure S4). Detailed characterization of clone #10 revealed a 13-bp deletion within the conserved loop of the ENE on the targeted allele. Notably, sequencing of the WT-sized upper band in this clone uncovered a collateral 1-bp thymidine (T) insertion within the ENE motif, suggesting that while the allele was not deleted, it likely underwent non-homologous end joining [30]. In the case of clone #31, sequencing confirmed a heterozygous state comprising one completely intact WT allele and one allele harboring a 45-bp deletion. This excision extended from the midpoint of U-rich motif 1 through the majority of U-rich motif 2, leaving only two distal nucleotides of the latter intact. As for clone #32, these cells harbored a 69-bp deletion including a segment upstream the ENE motif, the U-rich motif 1 and most of the conserved loop. Sequencing of the WT-sized band in clone #32 revealed a single nucleotide mutation (6892A>G) three nucleotides upstream the U-rich motif 1 (Figure 1F, Supplementary Figure S5). These results highlight the varying efficiency of CRISPR-Cas9 in MCF7 cells and provide a valuable model for comparing the effects of partial (clone #31) versus near-complete (clones #10, #32) structural disruption of the *MALAT1* 3’ end.

We next assessed the impact of these genomic modifications on *MALAT1* steady-state levels by RT-qPCR. In AGS cells, biallelic disruption of the 3’ ENE led to a highly significant reduction in *MALAT1* expression across all tested clones, with transcript levels falling to less than 10% of controls (Figure 1G; all *Padj*<0.0001). In the MCF7 model, clones #10 (harboring the deletion and collateral T-insertion) and #32 (with the deletion and the A-to-G mutation) exhibited a robust decrease in expression (*Padj*=0.0009 and *Padj*=0.0020, respectively). Interestingly, clone #31 (the heterozygous deletion) showed a non-significant reduction of approximately 25% (*Padj*=0.1389), suggesting that a single intact allele provides substantial levels of the transcript (Figure 1H).

To orthogonally validate these findings, we performed RNA-FISH using tiling probes targeting the full-length *MALAT1* transcript. In control cells, *MALAT1* exhibited its characteristic high-intensity nuclear signal (Figure 1I, J). Consistent with our RT-qPCR data, *MALAT1* signal was virtually abolished in the AGS biallelic clones (Figure 1I). In the MCF7 models, FISH signal was markedly attenuated in clones #10 and #32, while clone #31 showed a moderate decrease in signal intensity (Figure 1J).

### Loss of the 3’ Motifs Leads to Accelerated *MALAT1* Transcript Decay

While biophysical studies have established the triple-helix as a structural barrier against exonucleolytic decay [9], its requirement for *MALAT1* longevity within the endogenous cellular environment remains to be fully defined. To determine whether the reduced steady-state levels observed in our CRISPR-edited clones were a consequence of impaired RNA stability, we inhibited *de novo* transcription using Actinomycin D.

We first optimized the treatment conditions by performing a dose-response assay in AGS cells. After 12 hours of incubation, 1 μg/mL actinomycin D reduced *MALAT1* levels by approximately 50% (Supplementary Figure S6A). Notably, during these optimization steps, we observed that the Ct values of common reference genes, such as ***18S rRNA*** and ***GAPDH***, fluctuated by more than **2 cycles** following transcriptional inhibition, rendering them unsuitable for normalization (Supplementary Figure S6B). In contrast, *RPLP0* (Ribosomal Protein Lateral Stalk Subunit P0), which encodes a structural component of the 60S ribosomal subunit, exhibited more stability under these conditions (Supplementary Figure S6B). We further confirmed the higher stability of *RPLP0* relative to *18s rRNA* using digital PCR for absolute transcript quantification (Supplementary Figure S6C). Consequently, RPLP0 was utilized as the internal normalization control for all subsequent RNA stability experiments.

To quantify decay kinetics, we performed a time-course analysis, collecting RNA at 3, 6, and 10 hours post-Actinomycin D treatment. In control AGS cells, *MALAT1* displayed its characteristic “longevity” [31], with over 50% of the transcript remaining at the 10-hour mark (Figure 2A), confirming its stability in the presence of an intact 3’ ENE. In contrast, the decay rate was accelerated in all biallelically edited clones. Non-linear regression analysis using a one-phase exponential decay model revealed a drastic reduction in transcript half-life (t_1/2_) across the edited lines. Specifically, clone #1 exhibited a t_1/2_ of 4.9 hours, while clones #3 and #5 showed even lower stability, with half-lives of 1.4 hours and 2.4 hours, respectively (Figure 2A).

**Figure 2.**
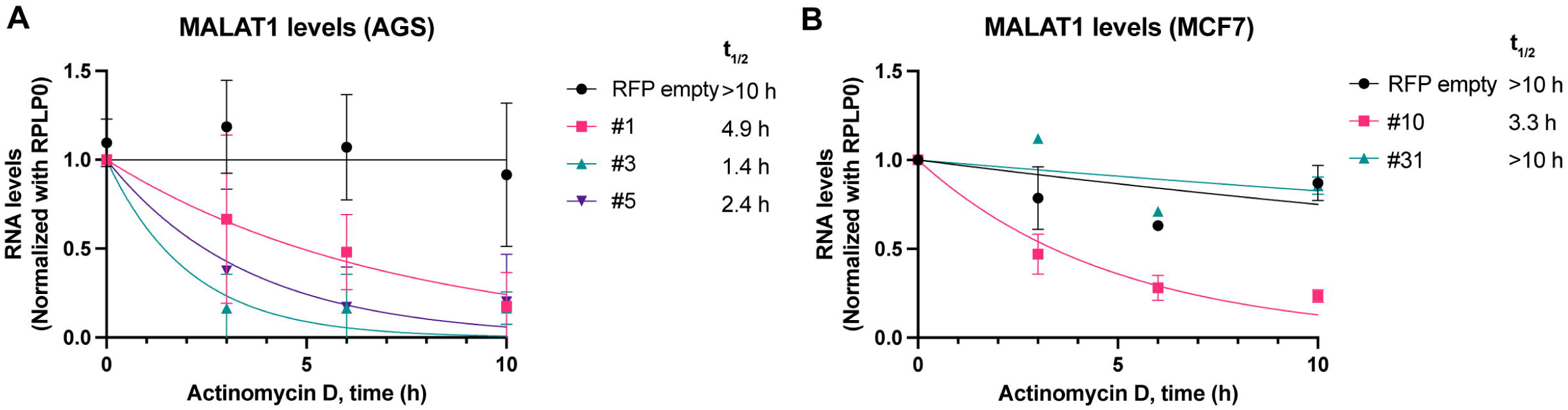
Disruption of the 3’ ENE accelerates *MALAT1* transcript decay. (A) RNA decay kinetics in AGS control and biallelic clones (#1, #3, #5) following 1 μg/mL Actinomycin D treatment times of treatment as shown. (B) RNA decay kinetics in MCF7 control and edited clones (#10, #31). Transcript half-lives (t_1/2_) were determined by fitting a non-linear one-phase exponential decay model. All data were normalized to the stable reference gene RPLP0. Results represent mean ± SD of three independent biological replicates.

In line with the previous results, control breast MCF7 cells exhibited a stable *MALAT1* profile over the 10-hour period (Figure 2B). In the edited MCF7 lines, clone #10 (harboring the deletion and collateral mutation) exhibited a t_1/2_ of 3.3 hours (Figure 2B). Interestingly, clone #31 (heterozygous deletion) maintained a stability profile similar to the control. Collectively, these results demonstrate that the 3’ triple-helix is determinant for the persistence of *MALAT1* in a cellular context.

### Deletion of *MALAT1* ENE produces a structured 3’ end

To determine whether the 3⍰-end deletion alters the structure of MALAT1, we performed in-cell DMS-MaP analysis of an amplicon spanning nucleotides 6335-7232, which directly encompasses the deleted 3⍰ ENE/triple-helix sequence. A previous study by Monroy-Eklund *et al*. [16] demonstrated that certain regions of MALAT1 exhibit strong structural conservation across cell types, experimental conditions, and even across species, despite differences in primary sequence. Based on these findings, we focused on the structured region spanning nucleotides 6537 to 6662 for comparative analysis in AGS cells, as this subdomain was identified as a region of low Shannon entropy, with a well folded structure [16]. Chemical modification of *MALAT1* performed in AGS clone #3 was successful, as DMS-treated samples exhibited higher per-nucleotide mutation rates than untreated control samples (Figure 3A), consistent with efficient probing of accessible A and C residues. DMS reactivities were highly reproducible between two independent biological replicates, with strong Pearson correlation (0.79) values supporting data robustness (Figure 3B). Structure modeling constrained by the average DMS reactivities from both replicates generated a secondary structure (Figure 3C) consistent with the previously reported MALAT1 folding architecture (Figure 3D). Highly reactive nucleotides mapped primarily to predicted single-stranded regions, whereas low-reactivity nucleotides localized to structured regions.

**Figure 3.**
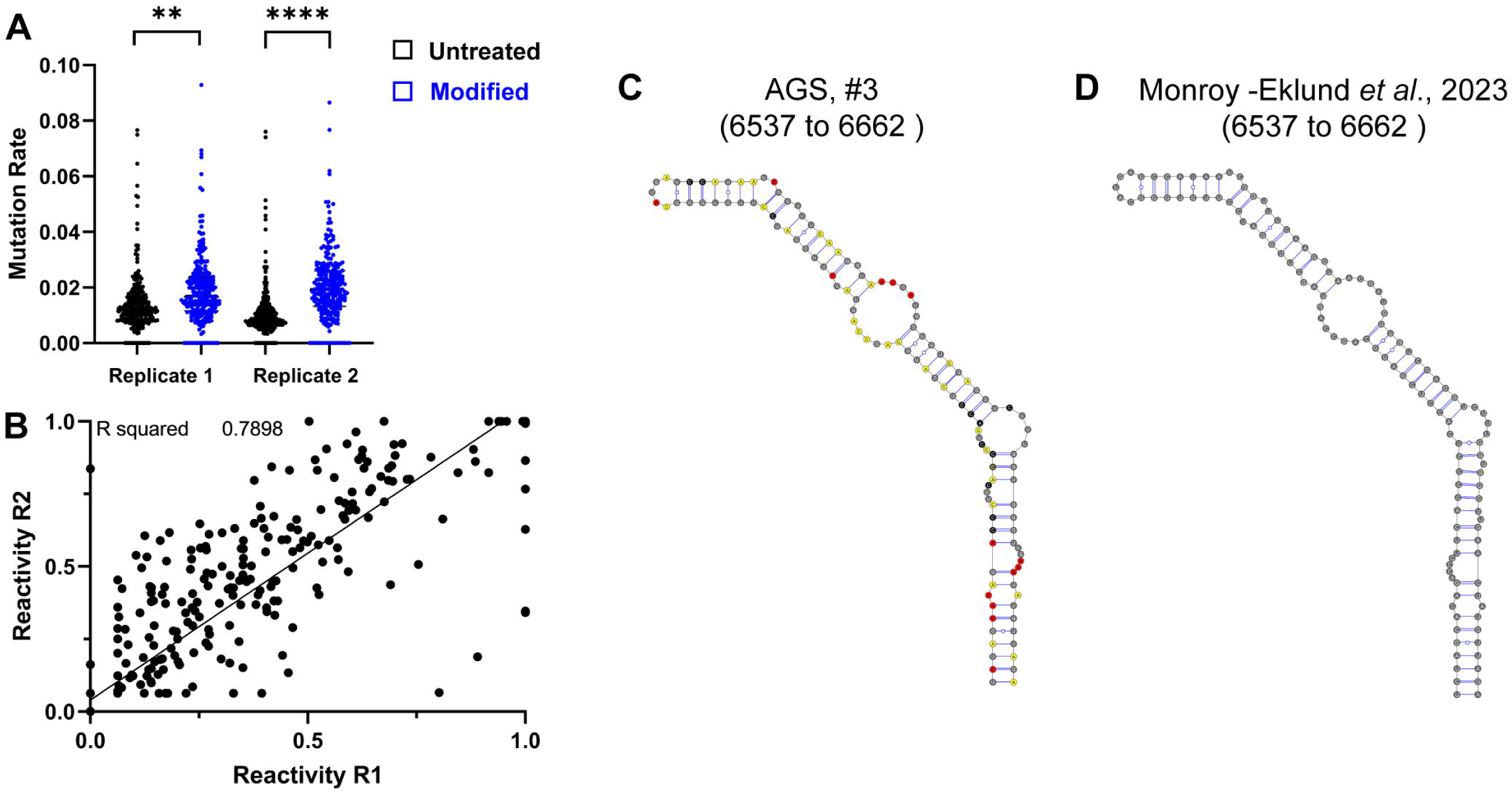
In-cell structural probing of MALAT1 with deletion in the 3’ end using DMS-MaP analysis. (A) Clone #3 of deleted AGS cells was chemically modified successfully. As expected, modified samples show higher mutation rates compared to control samples. Per-nucleotide mutation rates are displayed. (B) DMS reactivities were reproducible among different biological replicates. DMS reactivities at each nucleotide position are plotted and compared between the two biological replicates. Pearson correlation coefficient (r) values are indicated. (C) *In vivo* secondary structure model from this study (6537-6662) (average of two biological replicates). Nucleotides with high DMS reactivity are highlighted in red, nucleotides with medium DMS reactivity are highlighted in yellow, nucleotides with low DMS reactivity are highlighted in black. (D) Previously published in-cell secondary structure model of the MALAT1 region spanning nucleotides 6537-6662, adapted from the supplementary data of Monroy-Eklund et al. [16]

Comparing structures in high-Shannon-entropy regions is not meaningful, since these regions do not fold into a single well-defined structure, so we did not extend this comparison to the remainder of the probed amplicon. Reactivity and modeled structures for the extended regions close to the deletion in clone #3 (6626-6992, deletion starts at 6693 (Figure 1)) are provided in Supplementary Figure S7. Together, these findings suggest that the 3⍰-end deletion does not substantially alter this region’s structure.

### Genomic Disruption of the *MALAT1* ENE Motif does not alter mascRNA levels

Previous studies at the *MALAT1* locus have established the presence of the primary sense transcript (*MALAT1*), an antisense partner (*TALAM1*), and a processed, *MALAT1*-derived small RNA named mascRNA that is shuttled to the cytoplasm (Figure 4A) [1]. Because both *TALAM1* and mascRNA have been suggested to facilitate *MALAT1* 3’ end processing [15], [32] [12], we sought to determine if the structural disruption of the *MALAT1* ENE affects the levels of these associated transcripts.

**Figure 4.**
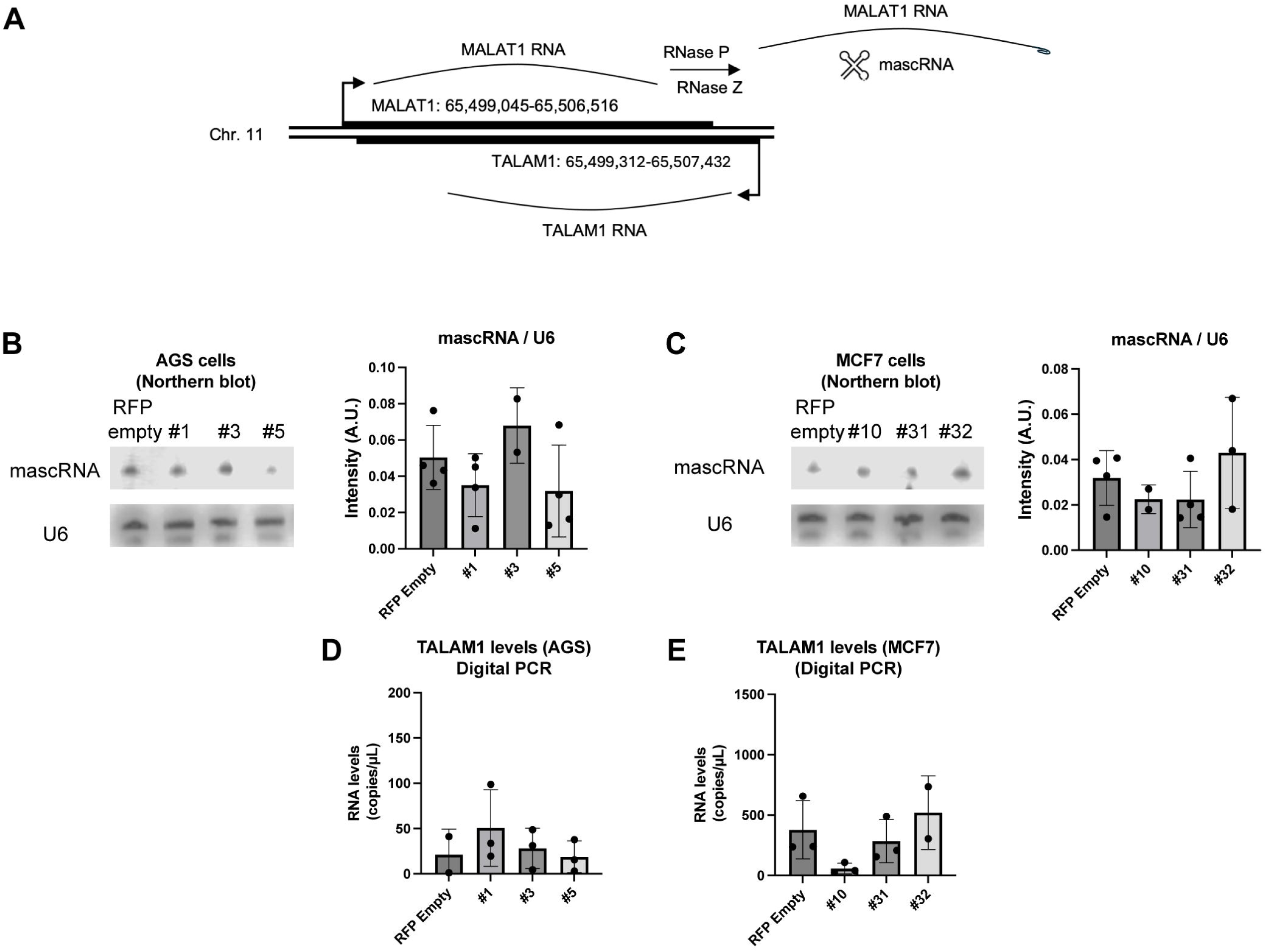
Impact of MALAT1 3’ end disruption on processed mascRNA and *TALAM1* levels. (A) Schematic of the coordinated expression of the MALAT1 locus, including the sense transcript, the antisense TALAM1, and the RNase P/Z-cleaved mascRNA. (B–C) Northern Blot analysis of mature, processed mascRNA in AGS (B) and MCF7 (C) clones. U6 snRNA was used as a loading control. Representative blots from at least two independent biological replicates are shown. Densitometric analyses were performed using Li-cor Image Studio v3.1.4. (D–E) Strand-specific RT-dPCR quantification of TALAM1 levels in AGS (D) and MCF7 (E) clones. Data represent mean ± SD from three biological replicates. Statistical comparisons across clones were evaluated by one-way ANOVA; no significant differences were observed (all *Padj* > 0.05).

First, we evaluated the expression of the processed mascRNA byproduct. Given its small size (58-61 nt [12]), we used Northern blot analysis to specifically detect the mature, processed form, rather than the primary transcript. Interestingly, despite the dramatic reduction in *MALAT1* levels shown in Figure 1, we observed no comparable changes in mature mascRNA expression in either the AGS (Figure 4B, all *Padj>0.05*; Supplementary Figure S8) or in the MCF7 edited clones (Figure 4C, all *Padj>0.05*; Supplementary Figure S8).

We next sought to measure the abundance of the antisense transcript *TALAM1*. To ensure accurate quantification and avoid signal interference from the high-abundance sense transcript, we performed reverse transcription with a tagged antisense-specific primer (Table 2) coupled with a digital PCR (dPCR) strategy designed to amplify tagged cDNA. Compared to *MALAT1* levels in control cells (17,600 copies/µL, CI (95%)=2.7% for AGS; 119,000 copies/µL, CI (95%)=2.7% for MCF7; Supplementary Figure S9A), baseline *TALAM1* abundance was markedly low in both AGS (21.25±28 copies/µL, Figure 4D) and MCF7 control cells (377±241 copies/µL, Figure 4E). Importantly, negative controls lacking reverse transcriptase or RT primers consistently yielded fewer than 10 copies/µL. Similarly low *TALAM1* levels were observed across all ENE-deleted clones (Figures 4D and 4E). To independently corroborate these findings, we analyzed Cap Analysis Gene Expression (CAGE) data from the FANTOM5 project. CAGE captures capped 5’ ends of RNA transcripts and is the gold-standard method for genome-wide identification of active transcription initiation sites [33, 34]. Analysis of both the complete FANTOM5 phase 2.5 dataset (Supplementary Figure S9B, “Total counts” track) and a cell-type-specific library for MCF7 cells (FANTOM5 Library ID: CNhs11943; Supplementary Figure S9B, “MCF7+” track), shows that the *MALAT1* TSS displays a sharp, well-defined plus-strand CAGE peak, confirming robust transcription initiation. In contrast, no localized minus-strand CAGE peak was observed at the annotated *TALAM1* TSS across any track (Supplementary Figure S9B, “Total counts” and “MCF7-” tracks). This absence is particularly evident in the magnified view (Supplementary Figure S9C) where the track scales confirm that the signal at the TALAM1 TSS is indistinguishable from background noise.

### Genetic Disruption of the *MALAT1* 3’ End Impairs AGS and MCF7 cell Proliferation

Finally, we investigated whether the destabilization of *MALAT1* translated into a quantifiable defect in cancer cell fitness. We monitored the growth kinetics of edited and control cells over a 72-hour period using automated cell counting, complementing with Annexin V/Propidium Iodide experiments to evaluate necrosis and apoptosis.

In the gastric cancer model, AGS clones harboring biallelic deletions exhibited a consistent and significant reduction in proliferation compared to control cells. The reduction in cell density became most pronounced toward the 72-hour mark, reflecting a sustained decrease in proliferative capacity following the loss of the *MALAT1* 3’ end (Figure 5A). Clones #1 and #5 displayed significant reductions (*Padj*=0.0171 and *Padj*=0.0026, respectively), while clone #3 exhibited the most robust phenotype at 72h (*Padj*<0.0001) (Figure 5A). Flow cytometry assays performed to evaluate the underlying cellular mechanism revealed that all three clones contained significantly higher proportions of apoptotic and necrotic cells compared to controls (Figure 5C). These results demonstrate that the levels of *MALAT1* transcript are determinants of survival in the analyzed AGS gastric cancer cells. Furthermore, the fact that all three independent biallelic clones showed impaired growth supports that the observed phenotype is a direct consequence of the ENE disruption rather than a clonal or off-target effect.

**Figure 5.**
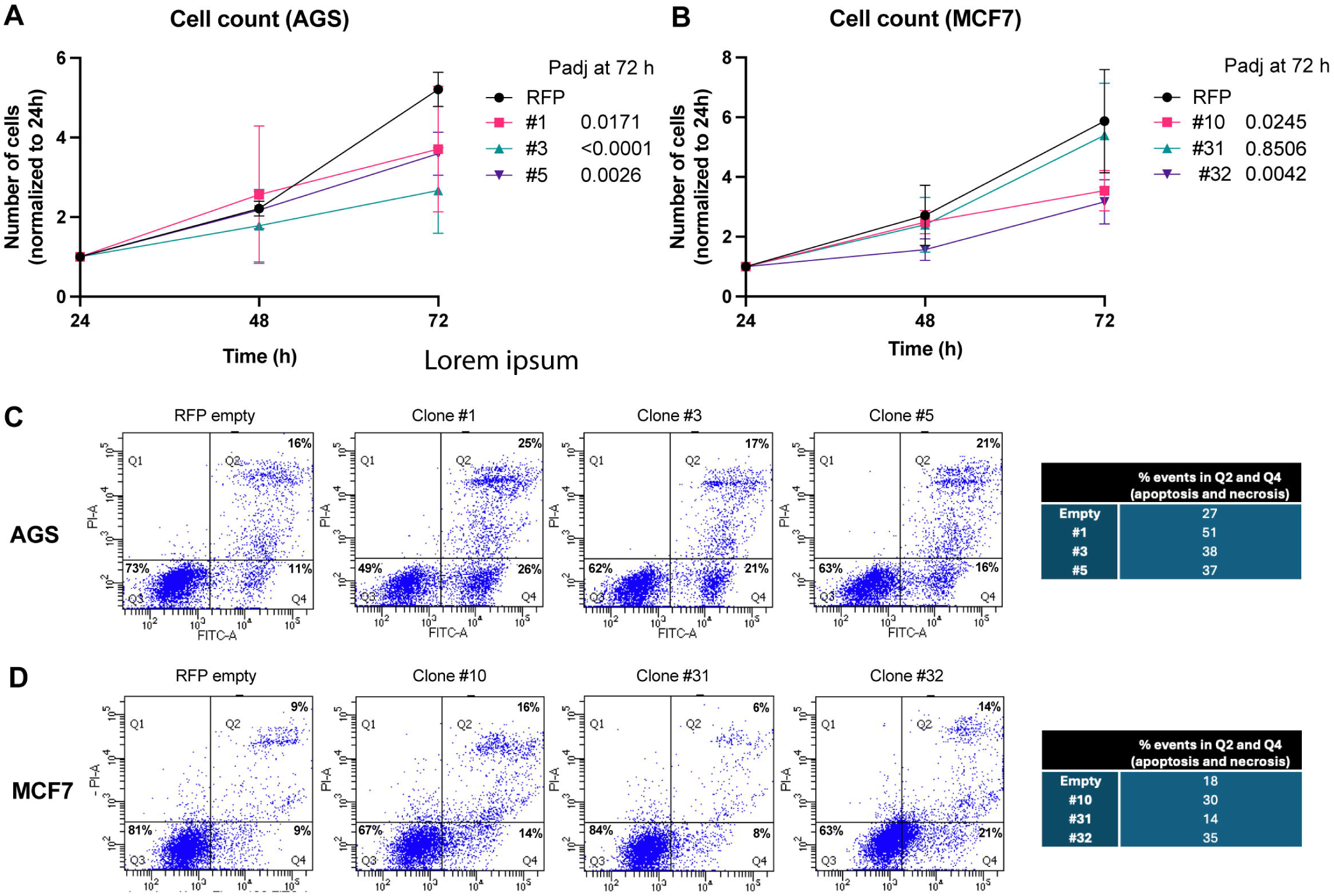
The MALAT1 3’ triple-helix is essential for proliferation in edited cells. (A) Growth curves of AGS control cells vs. biallelic clones #1, #3, and #5 over a 72 h period. (B) Growth curves of MCF7 control cells vs. edited clones #10, #31, and #32. Cell numbers were determined using a Luna-III automated cell counter with Trypan Blue exclusion for viability. Statistical significance versus controls was determined by two-way ANOVA. *Padj*-values for 72 h time points are indicated for each clone. Data are presented as mean ± SD from three independent experiments. (C-D) Flow cytometry analyses of apoptosis and necrosis in AGS (C) and MCF7 (D) cells, plotting Annexin V fluorescence (x-axis) against propidium iodide (PI) fluorescence (y-axis). One representative bivariate dot plots and quantitative summaries of total apoptotic cells (sum of quadrants Q2 + Q4) from three biological replicates are shown.

In the MCF7 breast cancer model, the reduction in cell proliferation and the induction of apoptotic and necrotic events mirrored the magnitude of *MALAT1* downregulation observed in our molecular assays. Clones #10 and #32, which exhibited the most dramatic loss of steady-state *MALAT1* levels, showed significant reductions in cell numbers at 72 hours (*Padj*=0.0245 and *Padj*=0.0042, respectively; Figure 5B) alongside the highest rate of cell death (Figure 5D). In contrast, clone #31 (the heterozygous deletion model with higher *MALAT1* levels) did not show changes with respect to control (*Padj*=0.8506) (Figure 5B) with apoptosis and necrosis rates remaining comparable to control levels (Figure 5D)

Collectively, these results indicate that the triple-helix-mediated stability of *MALAT1* is essential for maintaining the proliferation characteristic in the analyzed AGS and MCF7 cancer cells.

## Discussion

### The 3’ ENE as a strong contributor of *MALAT1* Metabolic Fate

Here we demonstrated that the genomic excision of triple helix-forming nucleotides from the *MALAT1* locus results in a profound loss of transcript stability (without affecting *MALAT1* byproduct mascRNA) and an overall decrease in the proliferative capacity of the analyzed gastric (AGS) and breast (MCF7) cancer cells linked to higher cell death. While previous efforts have utilized CRISPR-Cas9 to introduce single-point mutations in the proximity of the triple helix [18], our strategy involved deletions of varying lengths and positions within the ENE in the endogenous gene to determine the contribution of this region in the stability and cancer proliferation in live cells. Interestingly, we did not observe a correlation between the size of the genomic deletion and the severity of the transcript decay. Whether the deletion targeted the U-rich motifs or the conserved loop, the biological outcome remained consistent, likely triggering the collapse of the triple-helix architecture.

Our results in the MCF7 heterozygous clone (where single-base changes were sufficient to impair the transcript levels) further support the idea that even minor steric or sequence perturbations can disrupt the precise hydrogen-bonding network required for formation of the triplets, in line with previous reports [18]. Furthermore, as the effects were observed in two unrelated cancer cells lines (AGS and MCF7), our results indicate that the ENE-mediated protection strongly contributes to *MALAT1* longevity, rather than a cell-type-specific accessory. Notably, unlike in AGS cells, we were unable to isolate biallelic deletion clones in MCF7 cells. After screening 35 independent MCF7 clones (a line that also exhibited substantially higher baseline MALAT1 expression by digital PCR) we are tempted to speculate that MCF7 cells may be particularly sensitive to complete *MALAT1* loss, a hypothesis that will be tested in future studies.

Our findings align with ongoing therapeutic strategies aimed at downregulating MALAT1 by targeting its 3⍰ end using small molecules or antisense oligonucleotides [17, 35, 36]. Although our study examined a defined set of CRISPR-edited clones in MCF7 and AGS cells, these results expand upon the body of evidence established across diverse cell lines [7] and human tissue contexts [5]. Furthermore, our insights provide empirical support for emerging biotechnological applications that harness ENE motifs to engineer chimeric transcripts and RNA aptamers with enhanced cellular half-lives and stability[10, 11].

A critical question for us was whether the A-rich tract, which may mimic a poly(A) tail, could independently confer stability if the triple helix was disrupted. By specifically targeting the ENE while leaving the downstream A-rich tract intact, we have provided endogenous evidence that this tract, by itself, is insufficient for transcript protection. The A-rich tract alone, stripped of its protective structural pocket, cannot shield the 3’ terminus from exonucleolytic degradation. This reinforces the entwined nature of the triple helix as a singular, indispensable functional unit where the structure, rather than the sequence alone, dictates the metabolic fate of the RNA.

While our approach successfully knocked-out the *MALAT1* module, the resulting rapid decay of the transcript presents an inherent stability-function paradox. Because the triple helix is vital to the existence of the RNA, it is challenging to experimentally uncouple its role as a stabilizer from its potential role as a scaffold for protein interactions. For instance, studies have identified the triple helix as a binding site for the methyltransferase METTL6 [37] or the epigenetic silencer PRC2 [38, 39], conferring *MALAT1* an interesting role in RNA-mediated epitranscriptomic and epigenetic modifications. In our models, the drop in steady-state levels precluded us from investigating whether these protein-RNA interactions were disrupted due to the loss of the RNA molecule itself. Similarly, while *in vitro* simulations and SAXS analyses suggest that the ENE can adopt intermediate folding states [40] or be modified by epitranscriptomic marks like methyl-6-adenosine (m^6^A), pseudo uridine (Ψ), or m^1^Ψ [41], our data indicate that, in a living cancer cell, any deviation from the native 3’ architecture is a terminal event for the transcript. Importantly, we show that, although low in abundance, these edited transcripts remain structurally complex in the 3’ end.

### The impact of targeting the 3’-ENE over *MALAT1* processing

Our endogenous model provided us the opportunity of analyzing the fate of MALAT1-derived short transcript mascRNA. The persistence of mascRNA even in clones with the most severe MALAT1 depletion, highlights a significant mechanistic decoupling. Our results indicate that the initial transcription of the locus and subsequent processing by RNase P and RNAse Z are independent of the downstream decay of the primary *MALAT1* transcript. Thus, once *MALAT1* is transcribed, it is rapidly processed. This allows the mature mascRNA to persist in our stable cell lines, likely protected by its tRNA-like tertiary structure and cytoplasmic export [12]. Given the described role of mascRNA in promoting tumor metabolism in other contexts [42], it is notable that we observed growth defects despite the retention of mascRNA. This suggests that the downregulation of *MALAT1* is, on its own, sufficient to impair cancer cell fitness. However, it raises the intriguing possibility that a total locus knockout eliminating mascRNA together with long transcripts, could further exacerbate the anti-proliferative effect.

Finally, while previous studies established that *TALAM1* promotes *MALAT1* 3’ end folding and stability [15], two independent approaches (strand-specific RT-dPCR and CAGE analysis) could not detect *TALAM1* as an expressed transcript in our cell models. The convergence of these orthogonal datasets across our cancer models and across more than 1,800 human CAGE samples (Supplementary Figure S9, “Total count” track) substantially strengthens this conclusion. While we acknowledge that detecting low-abundance transcripts is inherently challenging and that context-specific expression under unexamined conditions or in specific tissues cannot be entirely ruled out, we found no empirical evidence for TALAM1 expression in our systems and were therefore unable to investigate its proposed regulatory mechanisms.

## Limitations of the study

While a consistent phenotype was observed across independent knockouts, sequence constraints inherent to targeting the 3⍰ triple-helix domain meant candidate sgRNAs possessed predicted off-target sites that were not experimentally evaluated. Furthermore, edited clones were not screened for gross genomic rearrangements. Consequently, we cannot definitively rule out off-target effects or clonal heterogeneity as drivers of the observed phenotype, which remains a limitation of this model.

## Data Availability Statement

The data underlying this article are available in the article and in its online supplementary material.

## Supplementary Data statement

Supplementary Data are available.

## Author Contributions Statement

Aramis Cortes-Arias, Valeria Valdes, Marcelo Muñoz-González, Diego Leiva, Andrew Acevedo, Michelle Tobar-Lara, Nicole Farfán, Lilly Oni, Eduardo Quiñones-Medina: Investigation, Visualization; Adriana Rojas, Roberto Munita, Srinivas Somarowthu, Veronica A. Burzio, Fernando J. Bustos: Writing – Review & Editing, Conceptualization, Methodology, Supervision, Formal analysis; Rodrigo Aguilar: Writing – Original Draft Preparation, Conceptualization, Methodology, Supervision, Formal analysis.

## Funding

ANID-FONDECYT 1230760 (VB), 11230662 (RM), 1250955 (FJB), 1240853 (RA); FONDEQUIP EQM230028 (FJB), Pew Advancement Award (RA, FJB)

## Conflict of Interest Disclosure

R.A. is an inventor on pending patent applications related to therapeutic strategies targeting *MALAT1*. The remaining authors declare no competing financial or personal interests.

## Declaration of generative AI and AI-assisted technologies in the manuscript preparation process

During the preparation of this work the author(s) used Gemini to assist with grammatical editing and language refinement. After using this tool, the authors reviewed and edited the content as needed and take full responsibility for the content of the published article.

## Acknowledgements

We thank the FACS-Sorting and digital PCR facility of the Institute of Biomedical Sciences at Universidad Andres Bello for its critical support during cell selection.

## Supplementary Figure Legends

**Supplementary Figure S1. Enrichment of RFP+ cells for CRISPR-Cas9 editing.** Representative FACS plots illustrating the selection and enrichment of RFP-positive cell populations 72 h post-transfection with CRISPR-Cas9 plasmids. These enriched populations were used for subsequent single-cell cloning and expansion of edited AGS and MCF7 lines.

**Supplementary Figure S2. Uncropped images of PCR screening and genotyping gels from AGS cells.**

**Supplementary Figure S3. Sanger sequencing validation of AGS clones.** Top: Schematic representation of the MALAT1 3’ end genomic sequence, highlighting the specific nucleotides excised in each biallelic clone. Bottom: Representative chromatograms showing the repair junctions for AGS clones #1, #3, and #5 compared to the wild-type (WT) sequence, confirming the successful deletion of the triple-helix-forming motifs.

**Supplementary Figure S4. Uncropped images of PCR screening and genotyping gels from MCF7cells.**

**Supplementary Figure S5. Sanger sequencing validation of MCF7 clones.** Top: Schematic representation of the monoallelic modifications in MCF7 clones. Bottom: Representative chromatograms for MCF7 clones #10, #31 and #32. For clone #10, the 13-bp deletion and collateral 1-bp (T) insertion are indicated. For clone #31, the 45-bp monoallelic deletion is confirmed. For clone #32, the 69-bp monoallelic deletion and the mutation is shown.

**Supplementary Figure S6. Optimization of Actinomycin D treatments and reference gene selection in AGS cells.** (A) Steady-state *MALAT1* levels measured by RT-qPCR after a 12 h incubation with varying concentrations of Actinomycin D to determine the optimal dose for stability assays. (B) Raw Ct values obtained by qPCR for three potential internal control genes (*GAPDH, 18s, and RPLP0*) under Actinomycin D treatment. (C) Absolute values (copies/µL) obtained by digital PCR for two potential internal control genes (*18s and RPLP0*) under Actinomycin D treatment *RPLP0* was selected for its superior stability across time points.

**Supplementary Figure S7. In-cell structural probing of MALAT1 with deletion in the 3’ end using DMS-MaP analysis (region closer to the ENE).** (A) MALAT1 Δ3’-3 (6626-6992) was chemically modified successfully. As expected, modified samples show higher mutation rates compared to control samples. Per-nucleotide mutation rates were displayed. (B) DMS reactivities were reproducible among different biological replicates. DMS reactivities at each nucleotide position were plotted and compared between the two biological replicates. Pearson correlation coefficient (r) values are indicated. (C) In vivo secondary structure model of MALAT1 Δ3’-3 (6626-6992) (average of two biological replicates). Nucleotides with high SHAPE reactivity are highlighted in red, nucleotides with medium SHAPE reactivity are highlighted in yellow, nucleotides with low SHAPE reactivity are highlighted in black

**Supplementary Figure S8. Uncropped images of Northern blot experiments.**

**Supplementary Figure S9. Complementary *MALAT1* and *TALAM1* level analyses.** (A) Absolute values (copies/µL) obtained by digital PCR for MALAT1 in control AGS and MCF7 cells (B) UCSC Genome Browser view of the *MALAT1*/*TALAM1* locus (chr11:65,264,209-65,275,903, hg19). *Tracks:* CAGE data from the FANTOM5 project showing transcription initiation signal. Total counts (aggregated across all FANTOM5 samples) and MCF7 cells are shown. The positive strand (*MALAT1* direction) is displayed in red and the negative strand (*TALAM1* direction) in blue. The vertical scale on the left of each track indicates the maximum signal intensity (counts or normalized values). The annotated TSS for MALAT1 and TALAM1 is shown. Note the absence of signal on the minus strand (*TALAM1* direction). (C) Magnified view of the annotated *TALAM1* TSS ± 200 bp showing the same CAGE data described in (B). Track scales confirm that the signal at the TALAM1 TSS is indistinguishable from background.

## Notes

### Summary of Updates

Expanded Experimental Models: Generation and characterization of an additional independent MCF7 mutant line (Clone 32), bringing our total dataset to 5 independent edited clones. Mechanistic Insights into Cell Survival: Annexin V and Propidium Iodide flow cytometry assays confirming that loss of MALAT1 stability triggers apoptotic and necrotic cell death. Target Specificity Controls: A targeted 895-bp upstream locus control deletion demonstrating that phenotypic outcomes are specific to ENE disruption rather than general genomic cutting at the locus. Absolute Transcript Quantification: Digital PCR (dPCR) absolute copy number quantification for MALAT1, TALAM1, and RNA decay reference controls (RPLP0). Orthogonal Validation & Nuanced Claims: Incorporation of FANTOM5 CAGE dataset analysis, moderation of universal claims, and transparent discussion of study limitations and off-target considerations.

