## Supplementary Figures for "Deletion of the *MALAT1* RNA 3’ end Promotes Transcript Decay and Inhibits Proliferation in Gastric and Breast Cancer Cells"

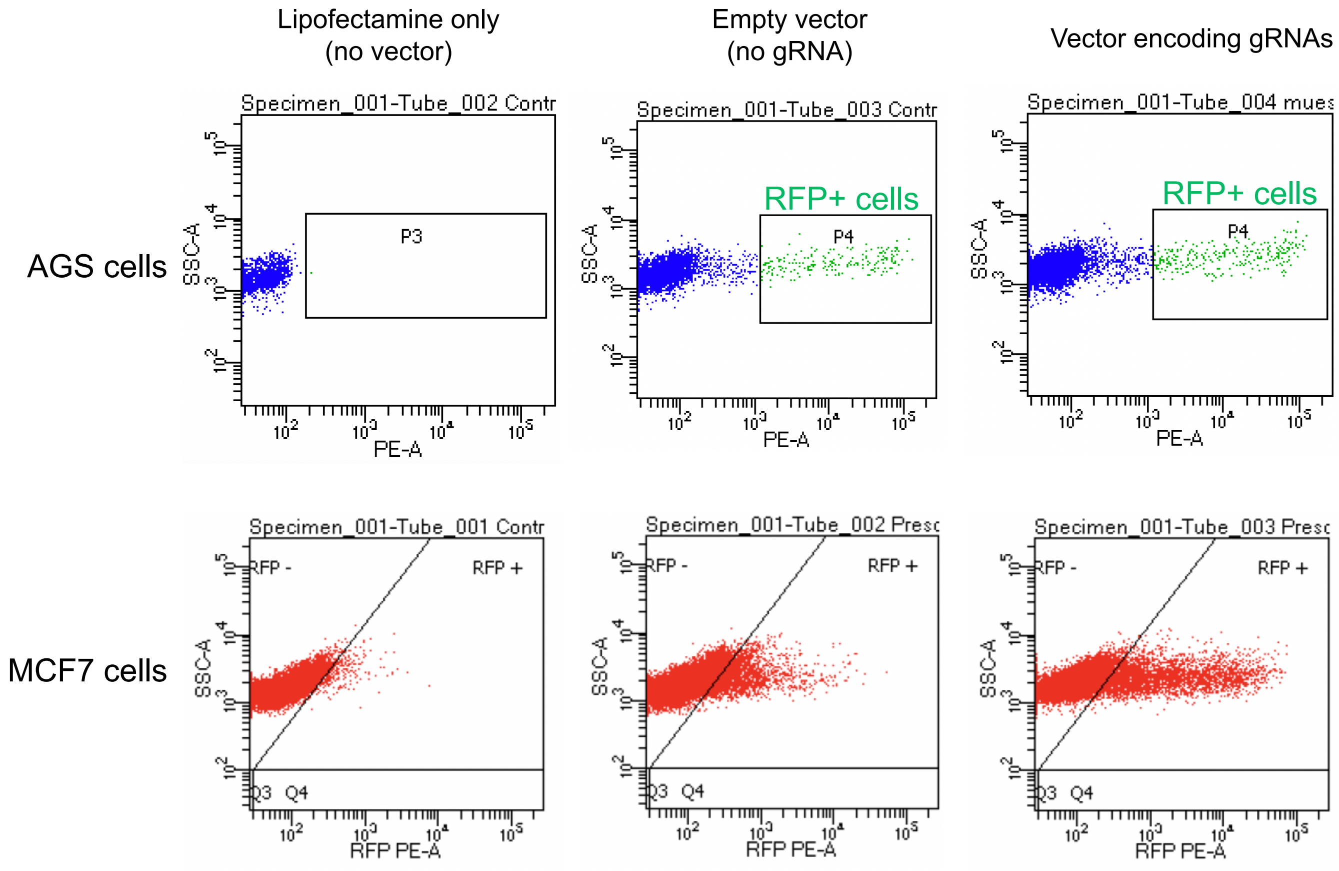


**Supplementary Figure S1**

### AGS cells

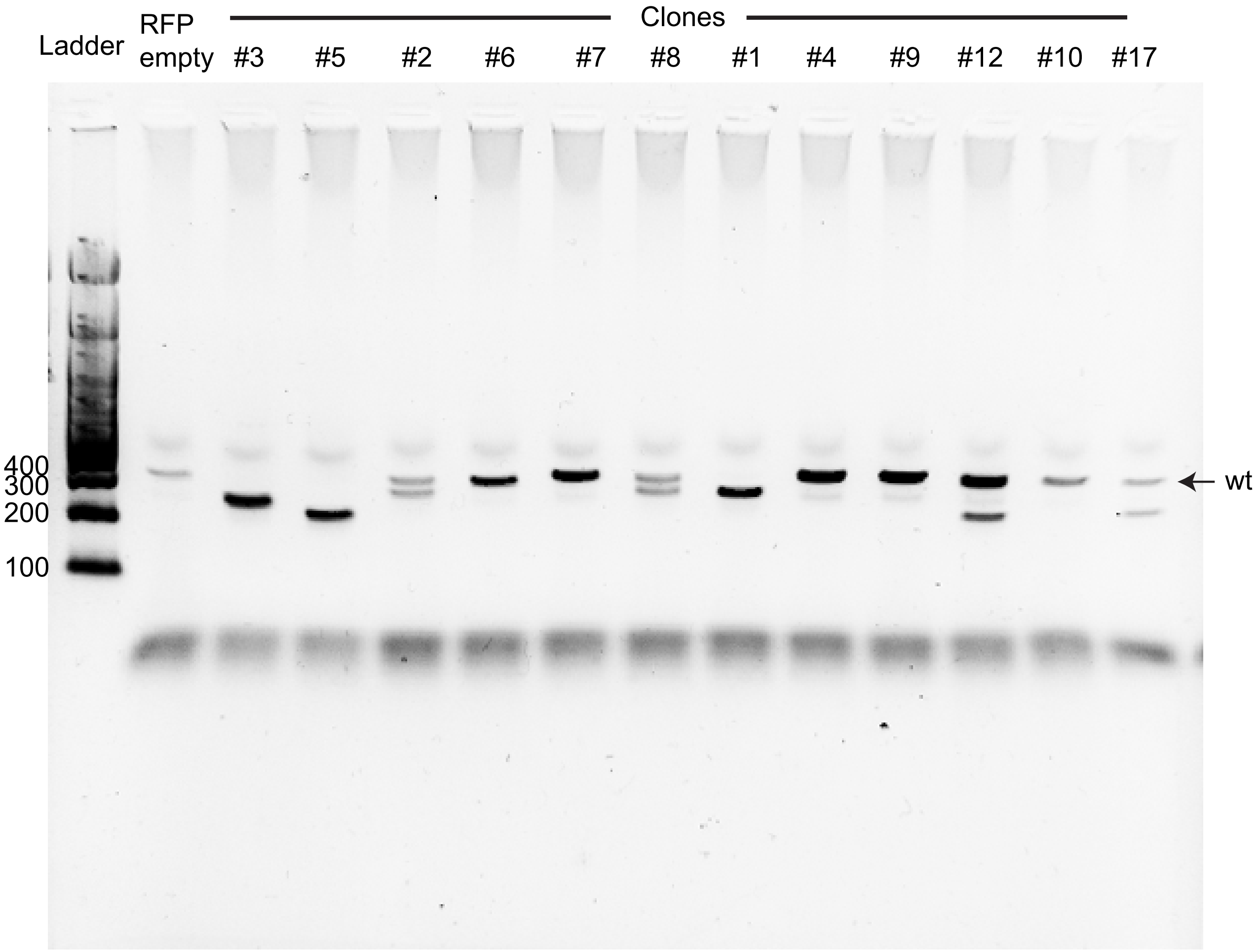


**Supplementary Figure S2**

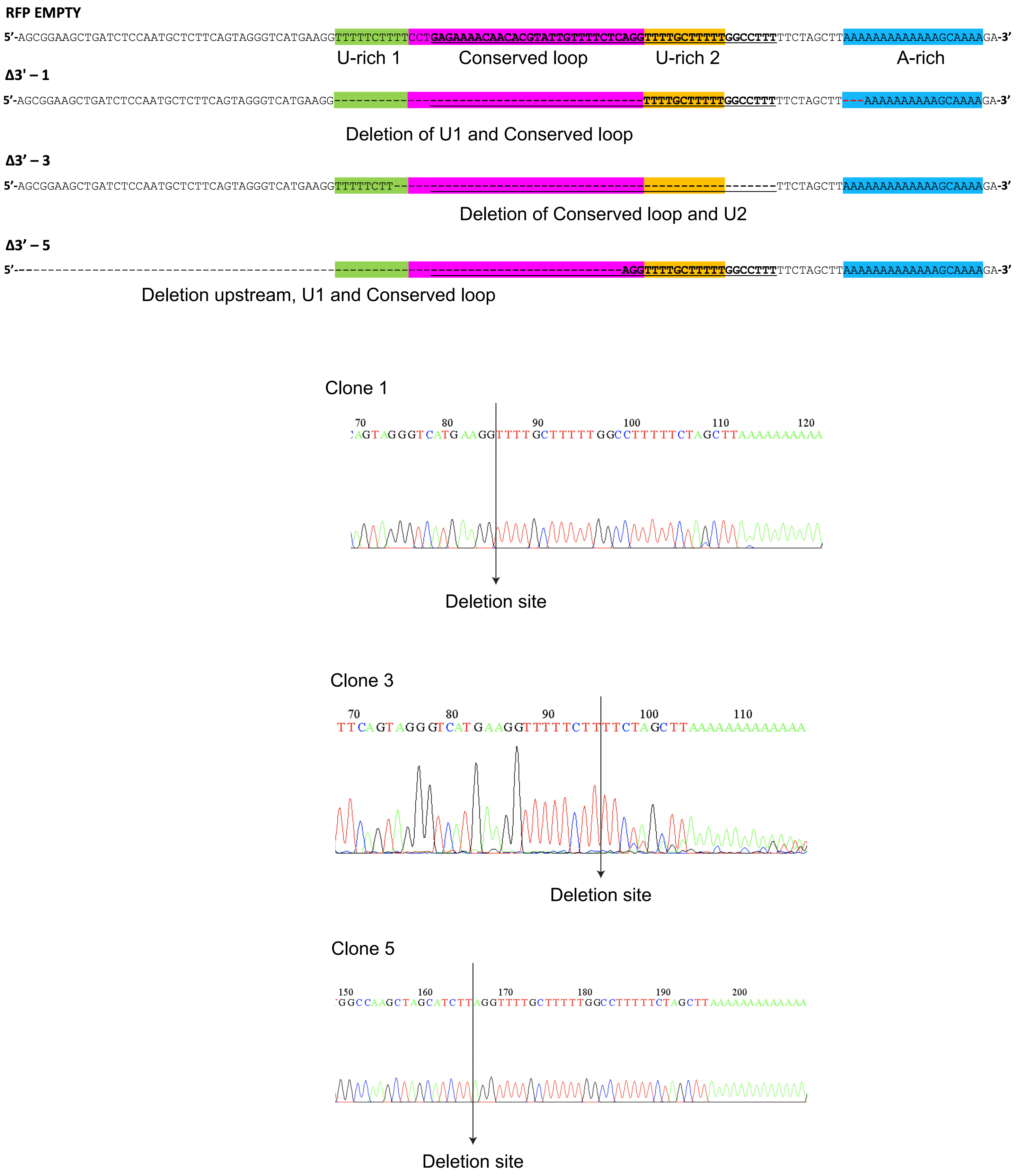


**Supplementary Figure S3**

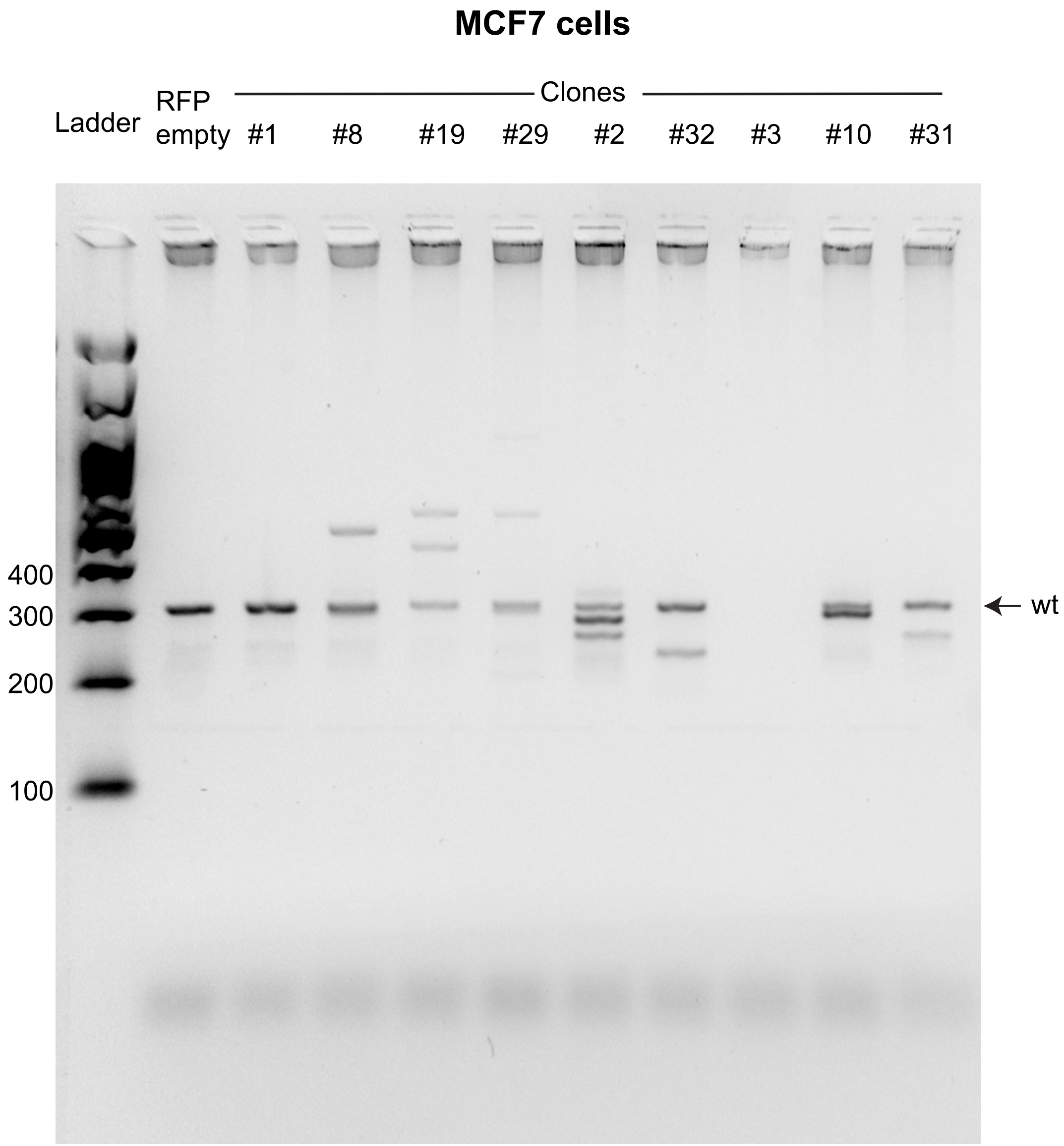


**Supplementary Figure S4**

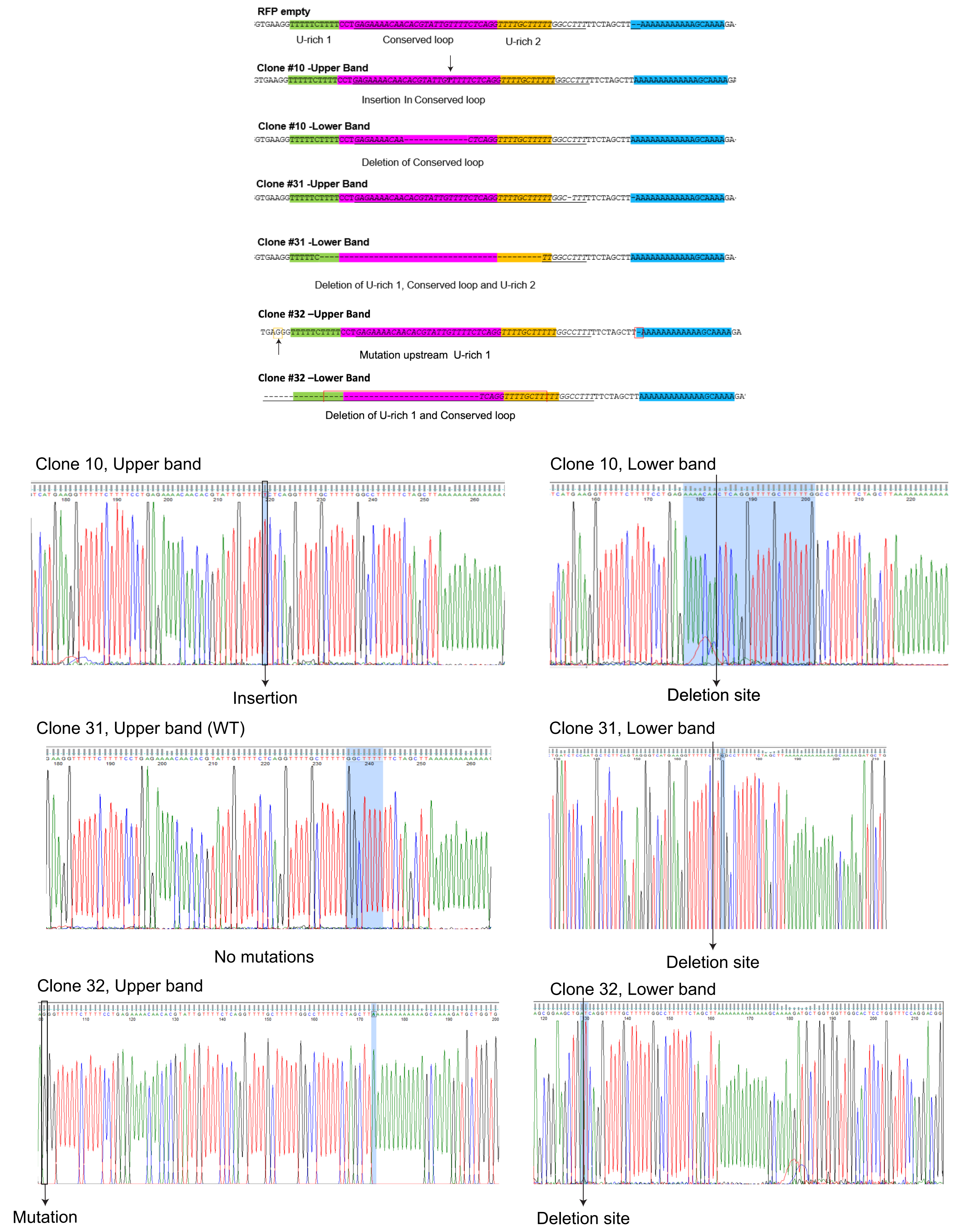


**Supplementary Figure S5**

A

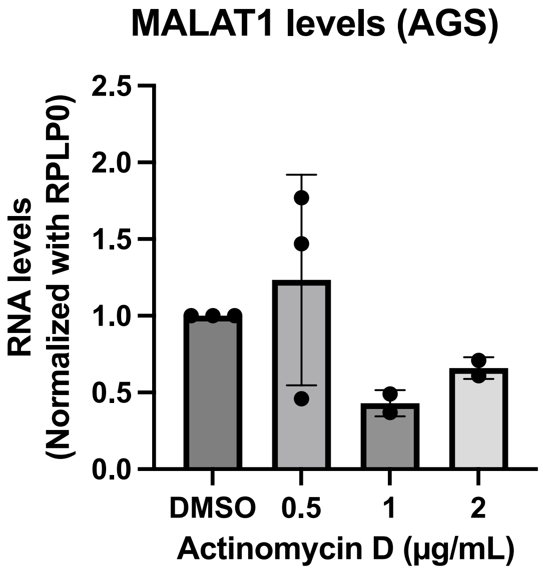


Housekeepking check

B, quantitative PCR (SYBR Green)

| | | | | |
|---|---|---|---|---|
| RPLP0 Ct | 16.16 | 17.03 | 16.76 | 16.83 |
| | 18.14 | 18.74 | 18.99 | 17.90 |
| | 17.90 | 18.52 | 19.89 | 18.15 |
| 18s Ct | 10.38 | 12.47 | 12.02 | 12.66 |
| | 10.58 | 12.95 | 13.08 | 12.27 |
| GAPDH Ct | 17.33 | 18.71 | 18.47 | 20.74 |
| | 17.95 | 18.79 | 20.54 | 18.61 |

Each row represents a different biological replicate

C, digital PCR

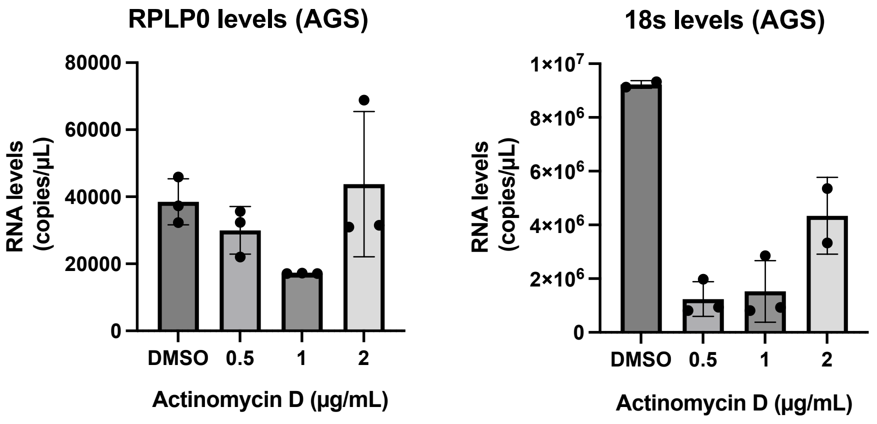


**Supplementary Figure S6**

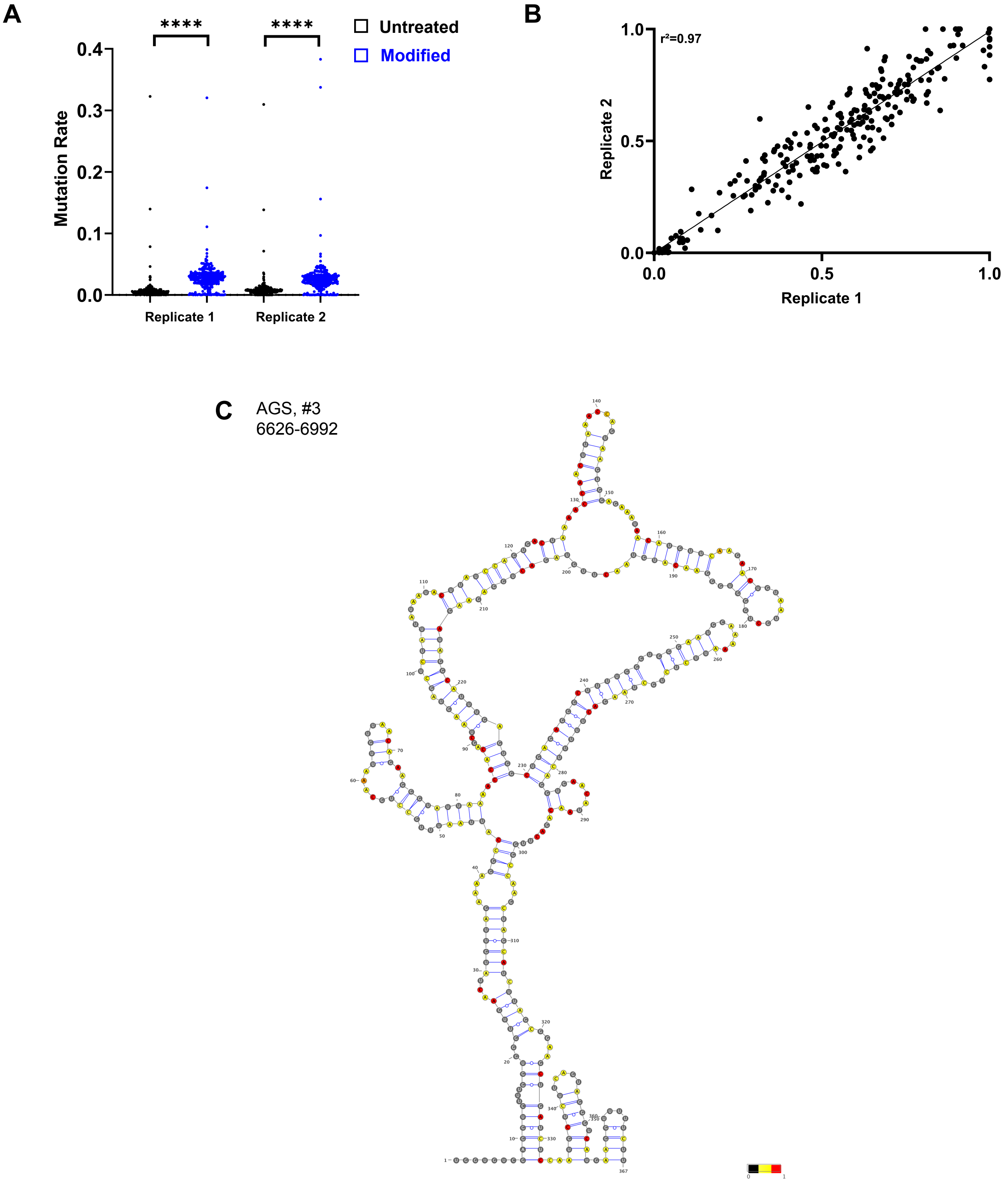


**Supplementary Figure S7**

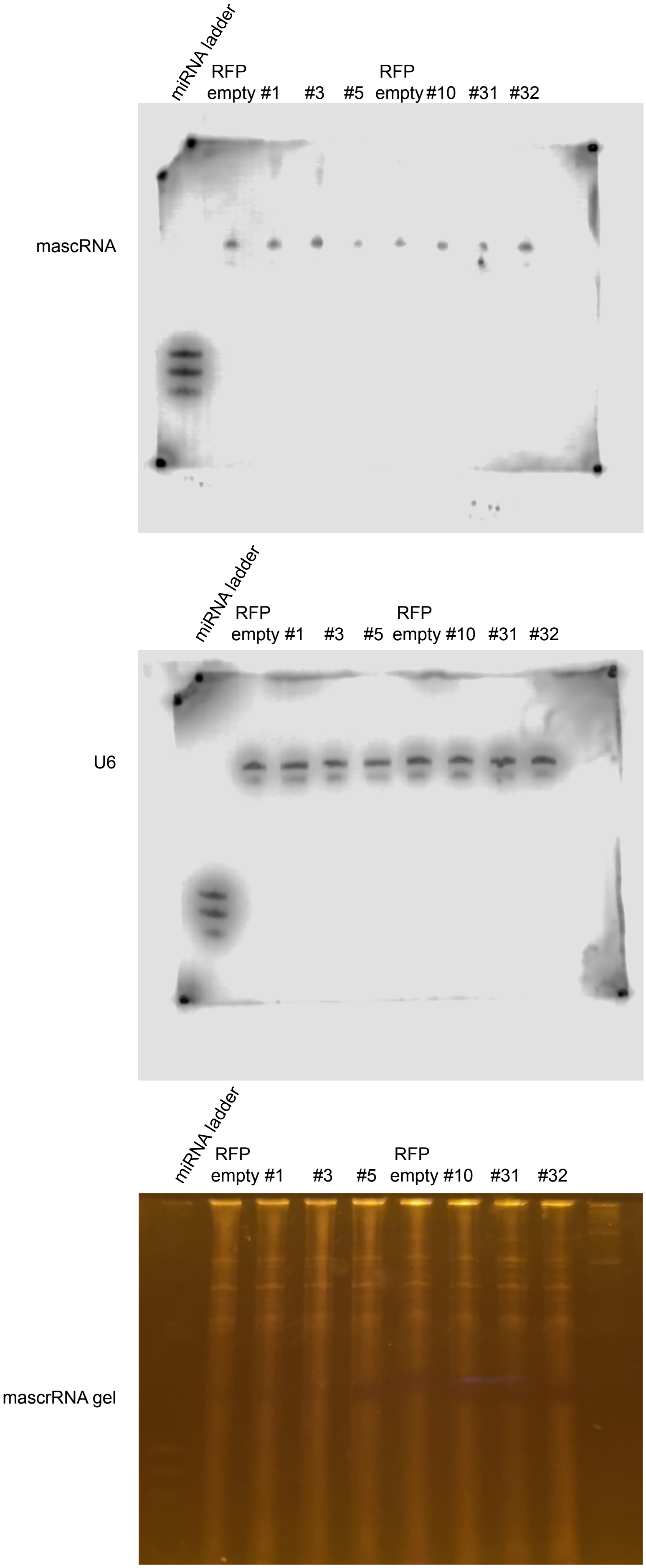


**Supplementary Figure S8**

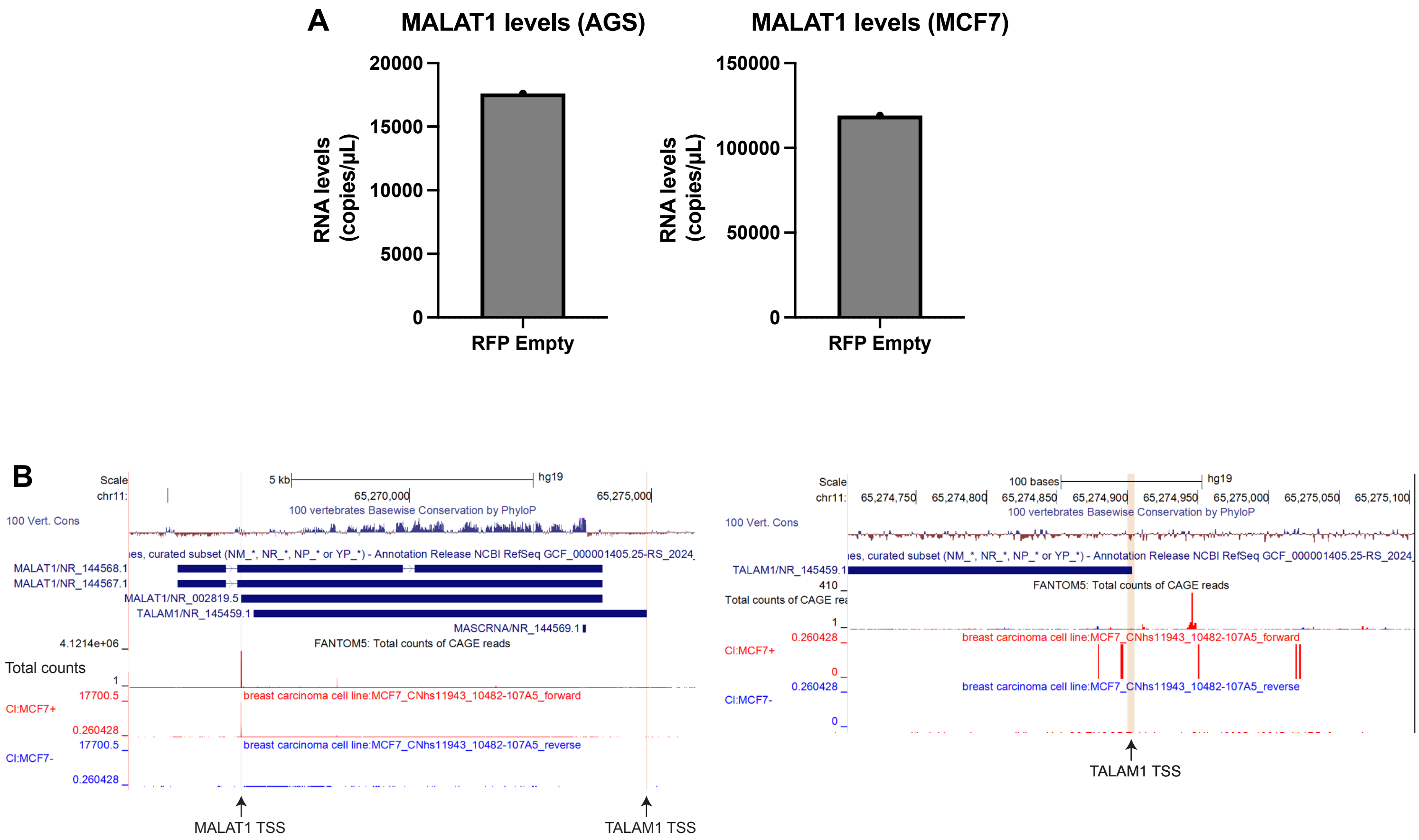


**Supplementary Figure S9**
